# Structures of two main components of the virophage and Marseilleviridae virions extend the range of unrelated viruses using fiber head as common receptor binding fold

**DOI:** 10.1101/2023.01.23.525297

**Authors:** Sandra Jeudy, Elsa Garcin, Alain Schmitt, Chantal Abergel

## Abstract

The detailed proteomic analysis of *Marseilleviridae* icosahedral capsids revealed that the two most abundant protein components of the virions were the Major Capsid Protein (MCP) and the product of an ORFan gene conserved in all *Marseilleviridae*. The noumeavirus NMV_189 3D structure revealed a common fold with fiber head proteins used by a variety of viruses to recognize their cellular receptor. However, the trimeric structure of NMV_189 uniquely lacking a tail domain, presented a deep concave site suggesting it could be directly anchored to the pseudo-hexagonal capsomers of the virion. This was confirmed by the unambiguous fit of the structure in the melbournevirus 4.4 Å cryo-EM map. In parallel, our structural genomic study of zamilon vitis virophage capsid proteins revealed that Zav_19 shared the same trimeric fiber head fold, but presented an N-terminal tail with a unique β-prism fold. The fiber head fold thus appears to be conserved in all types of non-enveloped icosahedral virions independently of their genomic contents (dsDNA, ssRNA, dsRNA). This could be a testimony of a common origin or the result of convergent evolution for receptor binding function.

**IMPORTANCE:** Giant viruses and their associated virophages exhibit a large proportion (≥60%) of orphan genes, *i.e*. genes without homologs in databases, and thus a vast majority of their proteins are of unknown function. The structural characterization of two ORFans, NMV_189 and Zav_19, both major components of noumeavirus and zamilon virophage capsids, respectively, revealed that despite a total lack of sequence homology, the two proteins share a common trimeric fold typical of viral receptor binding proteins and could be responsible for host receptor recognition. These two structures extend the range of unrelated viruses using fiber head structures as common receptor binding fold.

## INTRODUCTION

Since the discovery of marseillevirus, the first member of a family of large DNA viruses infecting Acanthamoeba (1), the *Marseilleviridae* family has been steadily expanding with now more than thirty members isolated from distant locations all around the world, among which 14 are fully sequenced (2). They seem to distribute among five lineages comprising marseillevirus (1), lausannevirus (3), tunisvirus (4), brazilian virus (5) and golden mussel (6) virus. They all present ~230 nm diameter icosahedral virions enclosing dsDNA genomes in the 340-400 kb range (1–9). The structures of melbournevirus and tokyovirus capsids have been determined (10, 11) and a model of the Major Capsid Protein (MCP) fitted into cryo-electron microscopy (cryo-EM) maps revealed an empty “cap” region on top of MCP. Proteomics analysis of melbournevirus and noumeavirus virions revealed that the most abundant protein was, with the Major Capsid Protein, an ORFan protein conserved in all *Marseilleviridae* (12). To understand the potential role of this abundant ORFan protein in capsid formation, we initiated the structural characterization of noumeavirus NMV_189. In parallel, proteomic analysis of the zamilon vitis virophage revealed that most (17/20) virophage-encoded proteins were present in the capsids. To gain some insights into their possible role, we initiated the structural characterization of ORFan proteins composing the capsids. Virophages of the *Lavidaviridae* family are 60 nm icosahedral viruses infecting giant viruses with cytoplasmic infectious cycle, diverting the giant virus viral factory to replicate. Because sputnik, the first discovered virophage, impairs the giant virus replication cycle, it was first proposed that virophages could in general protect host cells undergoing giant virus infections (13, 14). Such a protective role was later demonstrated for the protozoan *Cafeteria roenbergensis* infected by CroV, in presence of its virophage mavirus (15–17). It now appears that virophages can present different relationships with their viral host since zamilon appears to be commensal with the giant virus it is associated to (18–20). The structure of sputnik has been determined, as well as the MCP and penton proteins structures that were fitted into the cryo-EM density (21).

We report herein the 3D structures for noumeavirus ORFan protein NMV_189 and zamilon vitis ORFan protein Zav_19, which share unexpected structural features despite their lack of sequence homology. Structures revealed a typical trimeric fiber head fold that is found in adenoviruses fiber knobs, reoviruses attachment fiber head proteins, and with bacteriophages receptor binding domains (RBP). Such trimeric receptors appear to be used not only by viruses infecting prokaryotes and eukaryotes, but also by viruses infecting giant viruses. Our results on proteins from viruses as different as marseilleviruses and virophages suggest that attachment and cell entry by non-enveloped viruses occur by a common molecular mechanism, which could either be a consequence of distant evolutionary relationship or convergent evolution.

## RESULTS

### Structure determination of noumeavirus NMV_189

We had previously showed that besides the Major Capsid Protein (MCP), a 153 amino-acid protein was the most abundant protein in *Marseilleviridae* capsids (12). This protein has no homologue in public databases, and is thus an ORFan (22), but it is highly conserved within the *Marseilleviridae* family with ~80% protein sequence identity between noumeavirus NMV_189 and its most distant relatives. Recombinant NMV_189 protein was expressed in *E. coli*, purified, and yielded pure and soluble trimeric protein, after cleavage of the tag. We determined the structure by single wavelength anomalous dispersion (SAD) with a monoclinic crystal of the selenomethionylated protein. The resulting 1.9 Å resolution electron density map allowed building of all twelve molecules per asymmetric unit corresponding to four NMV_189 trimers. Data collection and refinement statistics are summarized in Table 1.

**Table 1.**
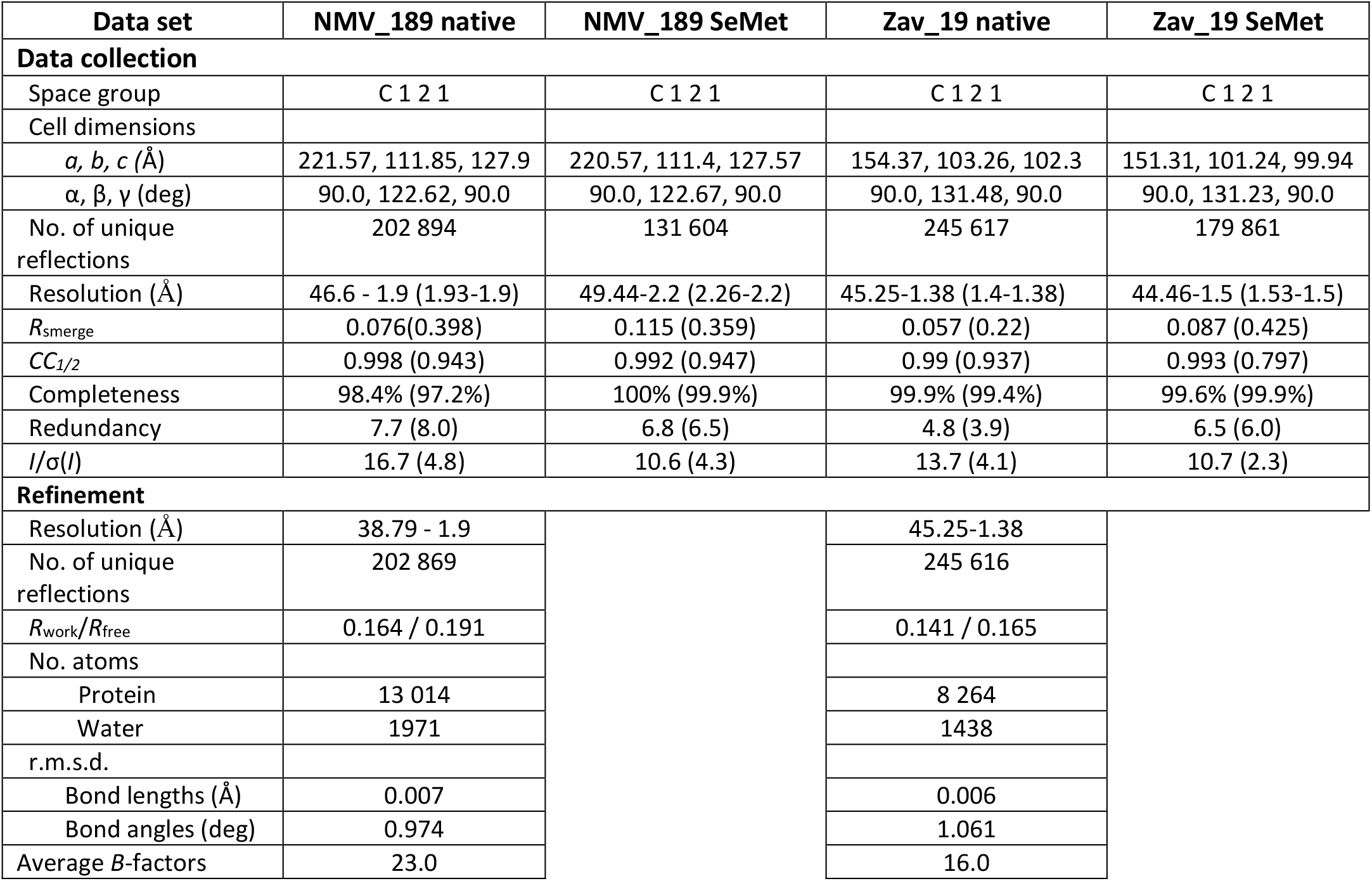
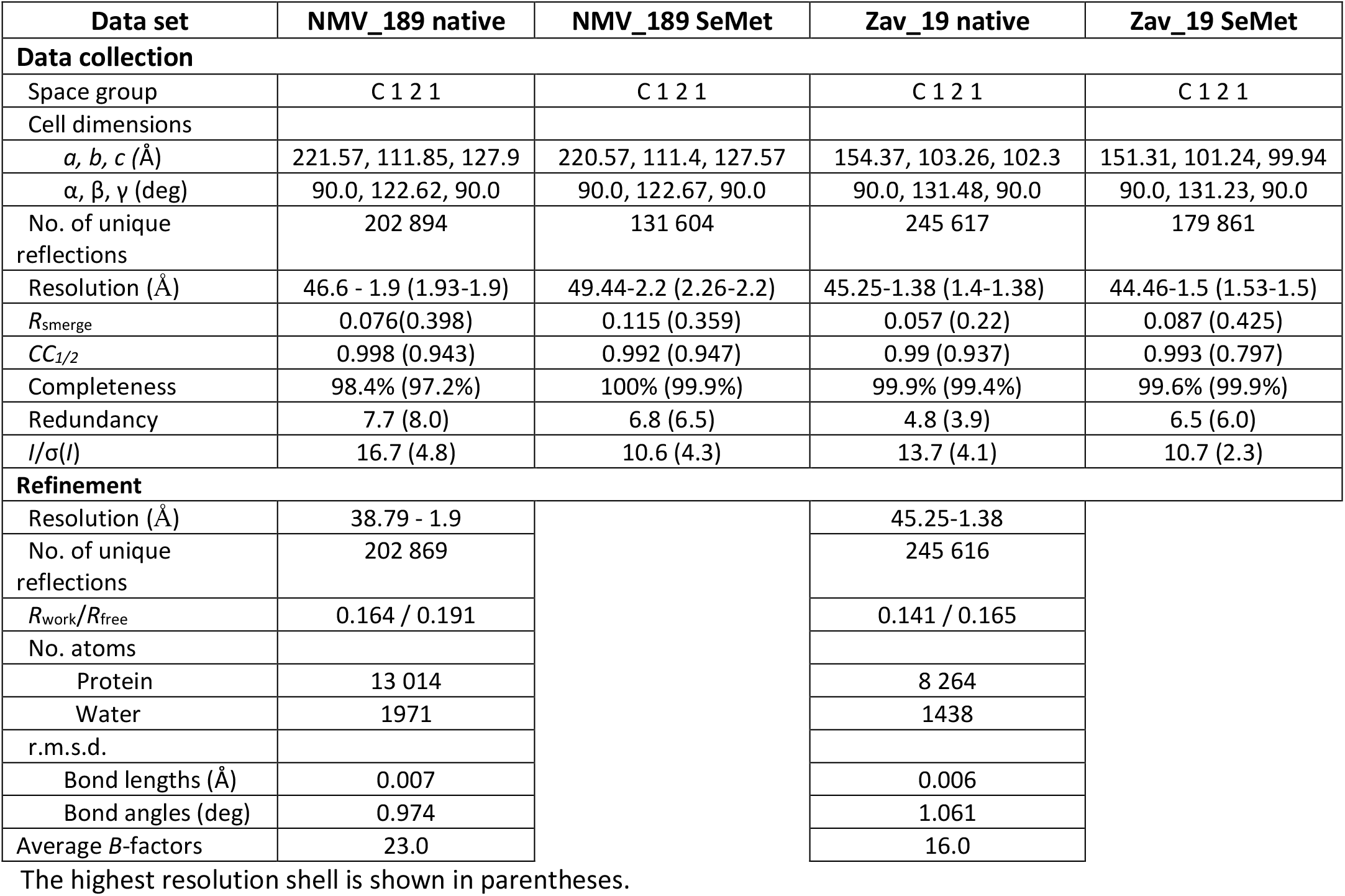
Data collection and refinement statistics

| Data set | NMV_189 native | NMV_189 SeMet | Zav_19 native | Zav_19 SeMet |
| --- | --- | --- | --- | --- |
| **Data collection** |  |  |  |  |
| Space group | C 1 2 1 | C 1 2 1 | C 1 2 1 | C 1 2 1 |
| Cell dimensions |  |  |  |  |
| $a, b, c$ (Å) | 221.57, 111.85, 127.9 | 220.57, 111.4, 127.57 | 154.37, 103.26, 102.3 | 151.31, 101.24, 99.94 |
| $\alpha, \beta, \gamma$ (deg) | 90.0, 122.62, 90.0 | 90.0, 122.67, 90.0 | 90.0, 131.48, 90.0 | 90.0, 131.23, 90.0 |
| No. of unique reflections | 202 894 | 131 604 | 245 617 | 179 861 |
| Resolution (Å) | 46.6 - 1.9 (1.93-1.9) | 49.44-2.2 (2.26-2.2) | 45.25-1.38 (1.4-1.38) | 44.46-1.5 (1.53-1.5) |
| $R_{\text{merge}}$ | 0.076(0.398) | 0.115 (0.359) | 0.057 (0.22) | 0.087 (0.425) |
| $CC_{1/2}$ | 0.998 (0.943) | 0.992 (0.947) | 0.99 (0.937) | 0.993 (0.797) |
| Completeness | 98.4% (97.2%) | 100% (99.9%) | 99.9% (99.4%) | 99.6% (99.9%) |
| Redundancy | 7.7 (8.0) | 6.8 (6.5) | 4.8 (3.9) | 6.5 (6.0) |
| $I/\sigma(I)$ | 16.7 (4.8) | 10.6 (4.3) | 13.7 (4.1) | 10.7 (2.3) |
| **Refinement** |  |  |  |  |
| Resolution (Å) | 38.79 - 1.9 |  | 45.25-1.38 |  |
| No. of unique reflections | 202 869 |  | 245 616 |  |
| $R_{\text{work}}/R_{\text{free}}$ | 0.164 / 0.191 | | 0.141 / 0.165 | |
| No. atoms |  |  |  |  |
| Protein | 13 014 |  | 8 264 |  |
| Water | 1971 |  | 1438 |  |
| r.m.s.d. |  |  |  |  |
| Bond lengths (Å) | 0.007 |  | 0.006 |  |
| Bond angles (deg) | 0.974 |  | 1.061 |  |
| Average $B$-factors | 23.0 | | 16.0 | |

The highest resolution shell is shown in parentheses.

Three NMV_189 monomers folded as an antiparallel beta sandwich of eleven beta strands (Fig. 1A) associate to form a trimer (Fig. 1B).

**Figure 1.**
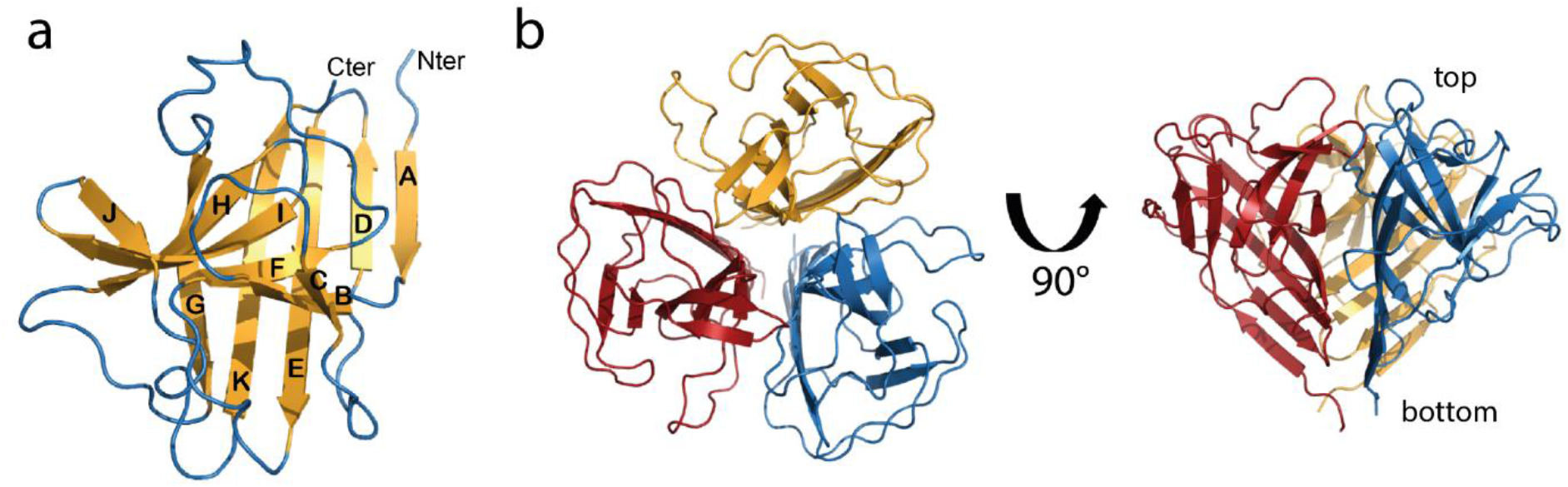
Ribbon representation of the NMV_189 structure. a] Nine out of the eleven beta strands form the two sheets of the beta sandwich. Strands A, D, E and K form the first sheet, and strands B, F, I and H form the second one, while the long G strand belongs to both sheets. Strands C and J are not involved in the sandwich although they belong to the second sheet. b] Orthogonal views of the color-coded monomers assembled into a trimer.

### Noumeavirus NMV_189 structurally resembles fiber head domains of other viruses

We used DALI (23) and VAST (24) servers to identify structural homologs and gain insights into NMV_189 protein function. The VAST search identified the closest structural homologs as the C-terminal head domain of the avian adenovirus CELO long fiber (PDB 2IUM (25), 2.7 Å rmsd for 95 residues out of 147) and type 1 short fiber (PDB 2VTW (26), 3.0 Å rmsd for 92 residues). The DALI search identified the closest homologue as the C-terminal domain of the tailspike protein of the salmonella phage Det7 (PDB 6F7K (27), with 2.7 Å rmsd for 123 residues). All these proteins are trimeric and are Receptor Binding Proteins (RBP) responsible for cell attachment, suggesting that NMV_189 could be involved in amoeba host cell attachment necessary to trigger phagocytosis. While adenovirus RBPs are made of three domains, including tail, shaft and head domains, NMV_189 only contains the globular head domain. This prompted us to hypothesize that it could be anchored to the capsid *via* direct binding to the Major Capsid Protein (MCP). Proteomic data analysis suggested a 1:1 stoichiometry (12), with one RBP trimer per trimeric MCP capsomer. Based on a model of noumeavirus MCP built from homologous PBCV-1 MCP structure (28), we showed that noumeavirus MCP capsomer and NMV_189 trimer display complementary surfaces and charge distribution (Fig. 2).

**Figure 2:**
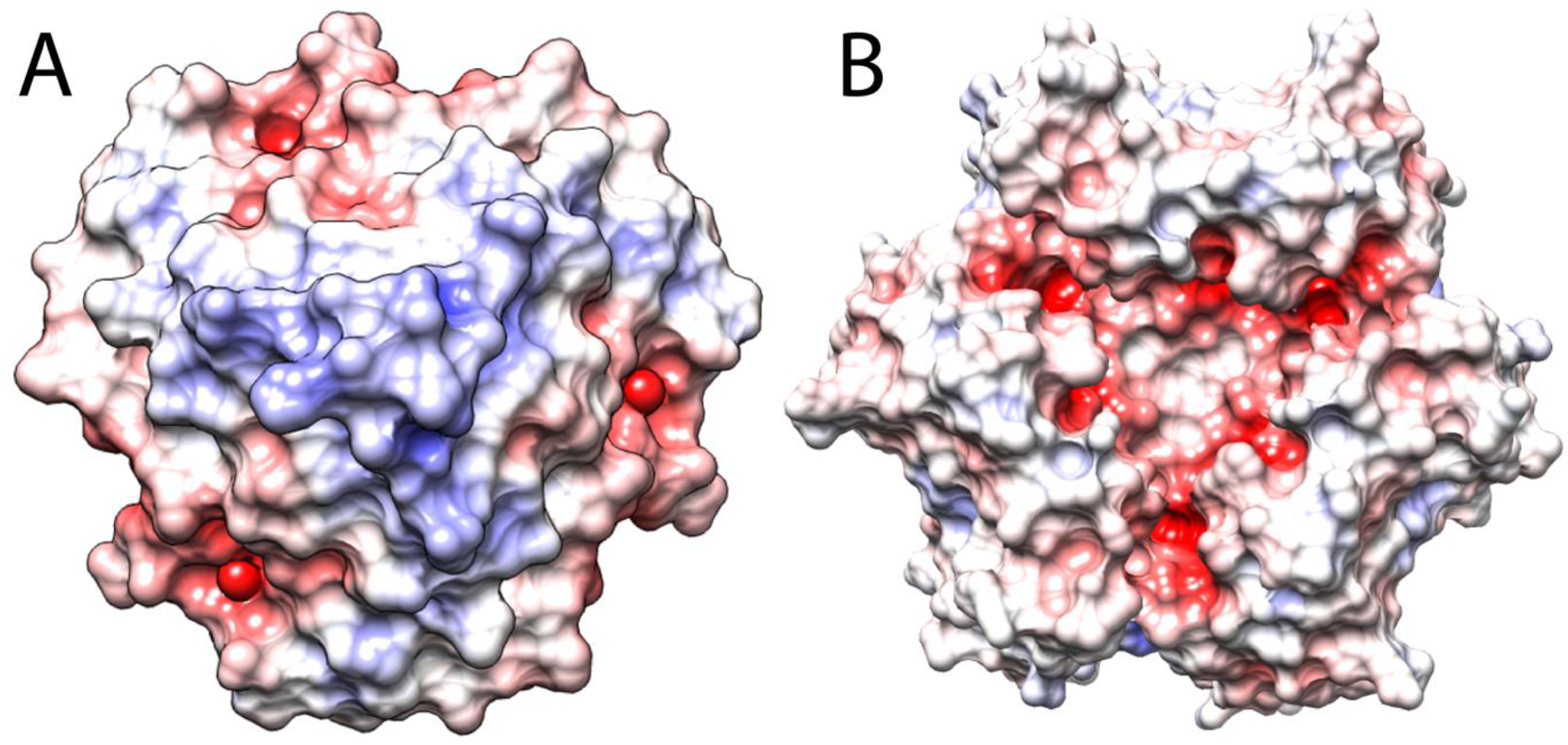
**Surface and charge complementarity** between A] NMV_189 trimer, bottom surface (see Fig.1) and B] noumeavirus capsomer model, top surface. Surfaces colored by electrostatic potentials calculated with APBS (29) ranging from red (–5 kT/e) to blue (+5 kT/e) were displayed with Chimera (30).

The tokyovirus capsid structure determined at 7.7 Å resolution (11) revealed an empty “cap” region on top of each MCP capsomer. Similarly, the 4.4 Å resolution structure (31) of melbournevirus particles confirmed an empty “cap” region above the MCP capsomers. Since NMV_189 shares 79% sequence identity with melbournevirus MEL_236, we fitted the NMV_189 trimeric structure in the empty region of the capsomer extracted from the cryo-EM map. The obtained model showed that trimeric NMV_189 could indeed fit above the trimeric MCP capsomer (Fig. 3) and that their interaction could be stabilized by electrostatic charge complementarity (see above).

**Figure 3:**
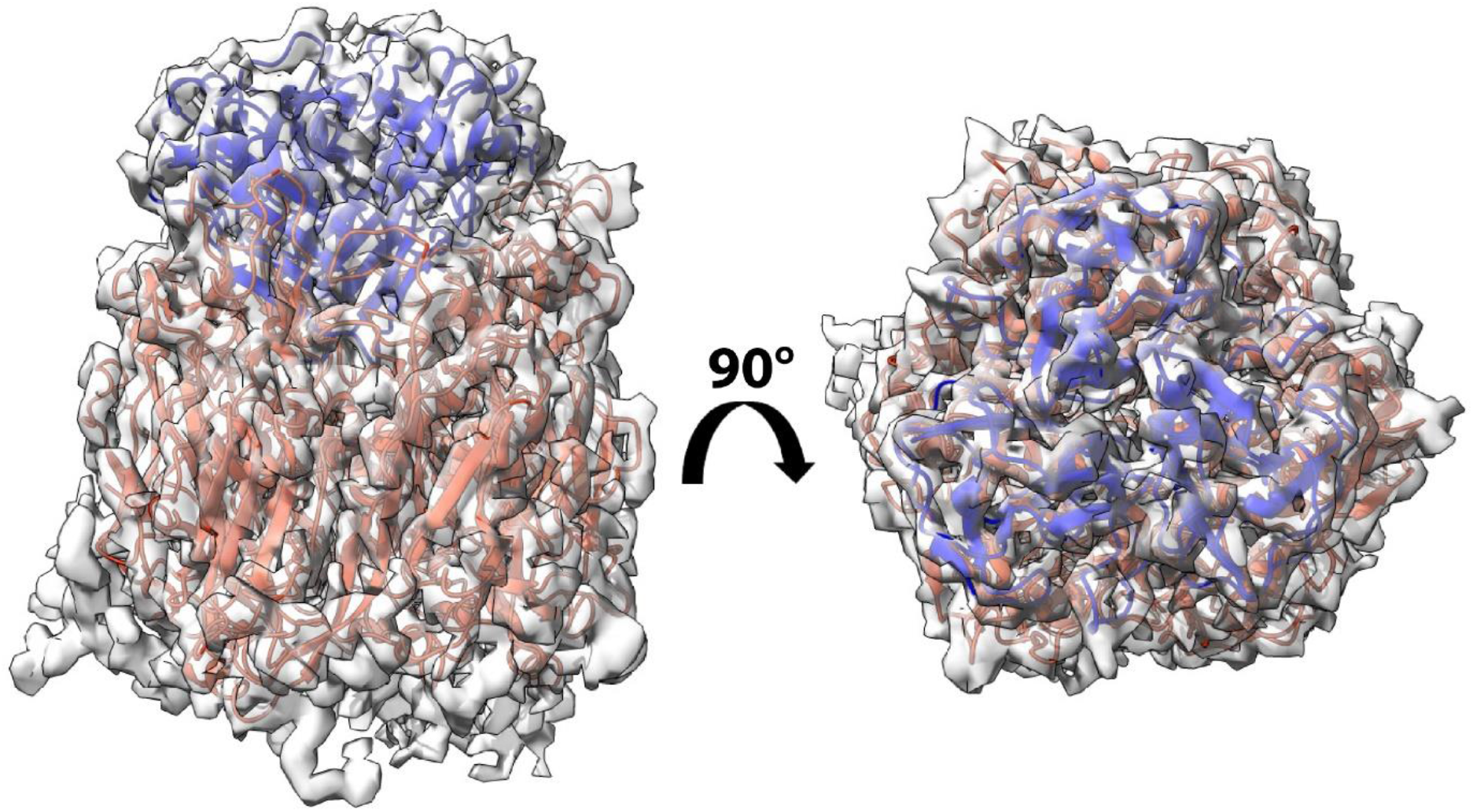
Fitting of melbournevirus MCP (in dark orange) and NMV_189 (in blue) into the segmented cryo-EM map of melbournevirus particle corresponding to a capsomer topped by a cap structure.

Interestingly, crystallographic electron density maps exhibited uninterpreted density localized in the central groove of each trimer (Fig. S1A). Serine residues (S109) from all three monomers surround the extra density and a layer of proline (P95) borders the cavity underneath the density while valine residues (V106) are exposed to the solvent on top of the head structure. We assigned the extra density to two cacodylate molecules present in the crystallization buffer in two different orientations (with one-half occupancy for each orientation). Using nano differential scanning fluorimetry (nanoDSF), we showed that sodium cacodylate increased the protein melting temperature (from 37.9°C to 41.6°C – Fig. S2), suggesting that NMV_189 binds cacodylate. Although both cacodylate molecules fit well in the density, peaks in the difference density map after refinement remained and therefore they were not included in the final structure.

### Structure determination of zamilon virophage Zav_19 protein

We cloned and expressed recombinant Zav_19 protein in *E. coli* cells. After tag cleavage, the 187-residue protein assembled as a soluble trimer. Crystals belonging to the monoclinic space group were obtained and the structure was determined by SAD using selenomethionine labeled-protein. There were six molecules per asymmetric unit corresponding to two trimers and the final model was refined to 1.38 Å resolution. Data collection and refinement statistics are summarized in Table 1.

Unlike NMV_189, Zav_19 is composed of two domains (Fig. 4): an N-terminal domain (residues 1-52) and a C-terminal domain (residues 61-184). The N-terminal domain folds into a unique β-prism structure composed of three beta-sheets (Fig. 4). These are made up of four strands that belong to all three monomers and that intertwine and pack around a pseudo three-fold axis. The N-terminal domains end with an 8 amino-acids α-helix that assemble into a three-helix bundle that connects the β-prism to the C-terminal domain (Fig. 4). The latter exhibits an antiparallel beta sandwich fold characteristic of viral fiber heads. Despite their lack of sequence identity, NMV_189 and Zav_19 trimers share a common fold and can be superimposed with a 3.3 Å rmsd for 304 residues (Fig. 5).

**Figure 4.**
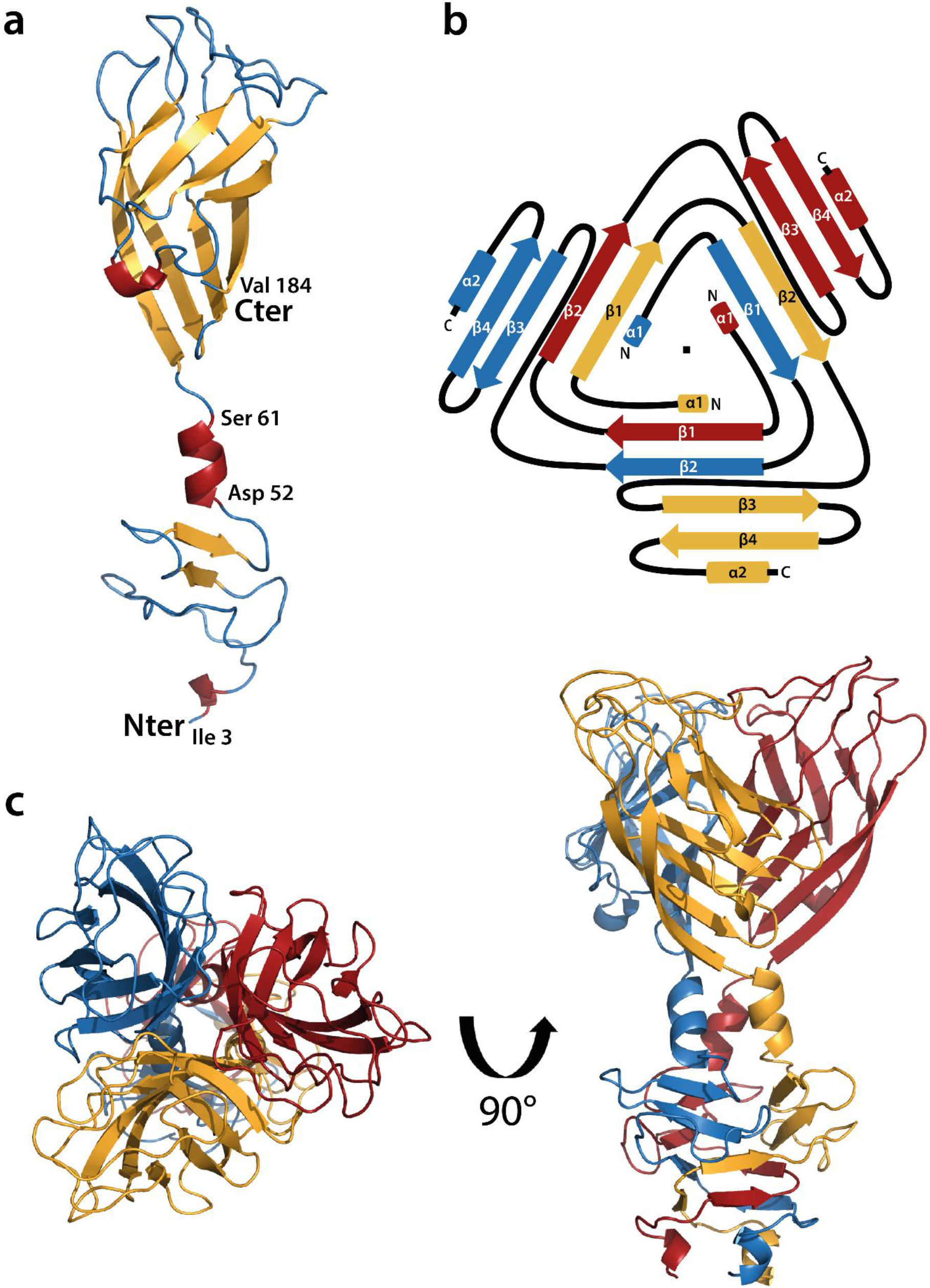
Ribbon representation of the Zav_19 protein structure. a] Monomer, b] Prism domain topology, the beta strand of each chain associates to the second strand of the adjacent molecule (parallel to the first one) and the third and fourth antiparallel beta strands of the third molecule of the trimer complete the 4 strands β-sheet. The three sheets pack around a pseudo three-fold axis and fold as a β-prism in the trimer c] trimer structure with a 3-helix bundle connecting the β-prism to the fiber head.

**Figure 5:**
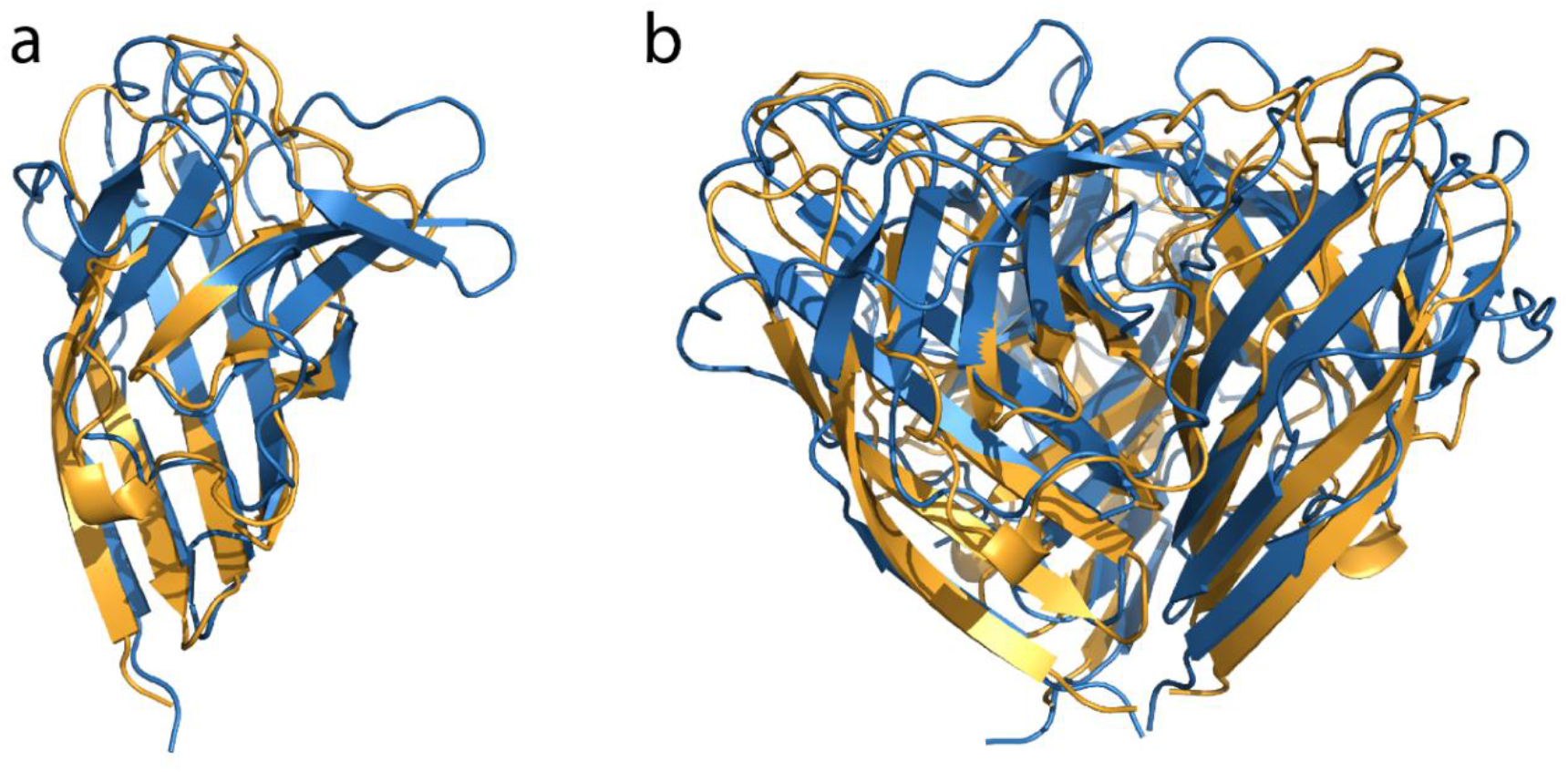
Superimposition of the a) monomeric and b) trimeric structures of NMV_189 (blue) and Zav_19 C-terminal domain (light orange). The two trimeric structures exhibit a 3.3 Å rmsd over 304 residues based on Cα superimposition.

We searched the PDB for Zav_19 closest structural homologues using Dali (23) and Vast (24) servers with the full length structure and each of the two Zav_19 domains independently. The C-terminal domain resembles viral receptor binding proteins, with one of the closest homolog being the RBP from Lactococcal phage TP901.1 (BppL protein, PDB 3EJC) (32). The C-terminal domain of the phage RBP monomer (residue 63 to 164) can be superimposed to Zav_19 C-terminal head domain (residues 61 to 184) with 2.5 Å rmsd (for 91 aligned residues out of 102, with 17% sequence identity). The trimeric C-terminal structures superimpose well with a 3.1 Å rmsd. The phage protein is part of the baseplate and is responsible for the specific recognition of saccharidic receptors localized on the host cell surface.

The conical N-terminal domain does not exhibit clear homology with other structures in the databases except for a β-prism fold (Fig. 4). Each monomer provides four β-strands that interlace to form the prism (Fig. 4B). The β-prism fold is also associated to various structures of lactococcal phages RBP (phage TP901-1: PDB 2F0C (33); phage P2: PDB 2BSD (34); phage Tuc2009: PDB 5E7T (35)) but it corresponds to the neck domain that connects the N-terminal to the C-terminal head domain. In the Zav_19 structure, the diameter of the β-prism is larger due to the long loop connecting strands β2 to β3 from each monomer that protrude outside of the β-prism.

Zav_19 shares 70% identity with its orthologous protein V8 in sputnik virophage. Given the close relationship between the two virophages, we looked at the cryo-EM reconstruction of sputnik. Interestingly, at 10.7 Å resolution, the Cryo-EM map revealed a “mushroom”-like fiber with a triangular head that appeared to protrude from the center of the pseudo-hexameric capsomer (36). Since these fibers were not visible in the 3.5 Å resolution reconstruction of the Sputnik particle (21), the authors concluded that they were likely flexible. It is tempting to attribute this extra density to the V8 protein in sputnik, orthologous to Zav_19 in the zamilon particle. The surface and charges of Zav_19 are complementary to zamilon MCP model (Fig. 6). However, Zav_19 is only the 7th most abundant protein in the capsids (Supplementary Table 1) with an estimated 50 copies per particle, a number that is incompatible with a one-to-one stoichiometry with MCP, thus preventing capsomers from being fully occupied. Alternatively, Zav_19 could have been lost during the 3-steps purification of the virophage capsids (36).

**Figure 6:**
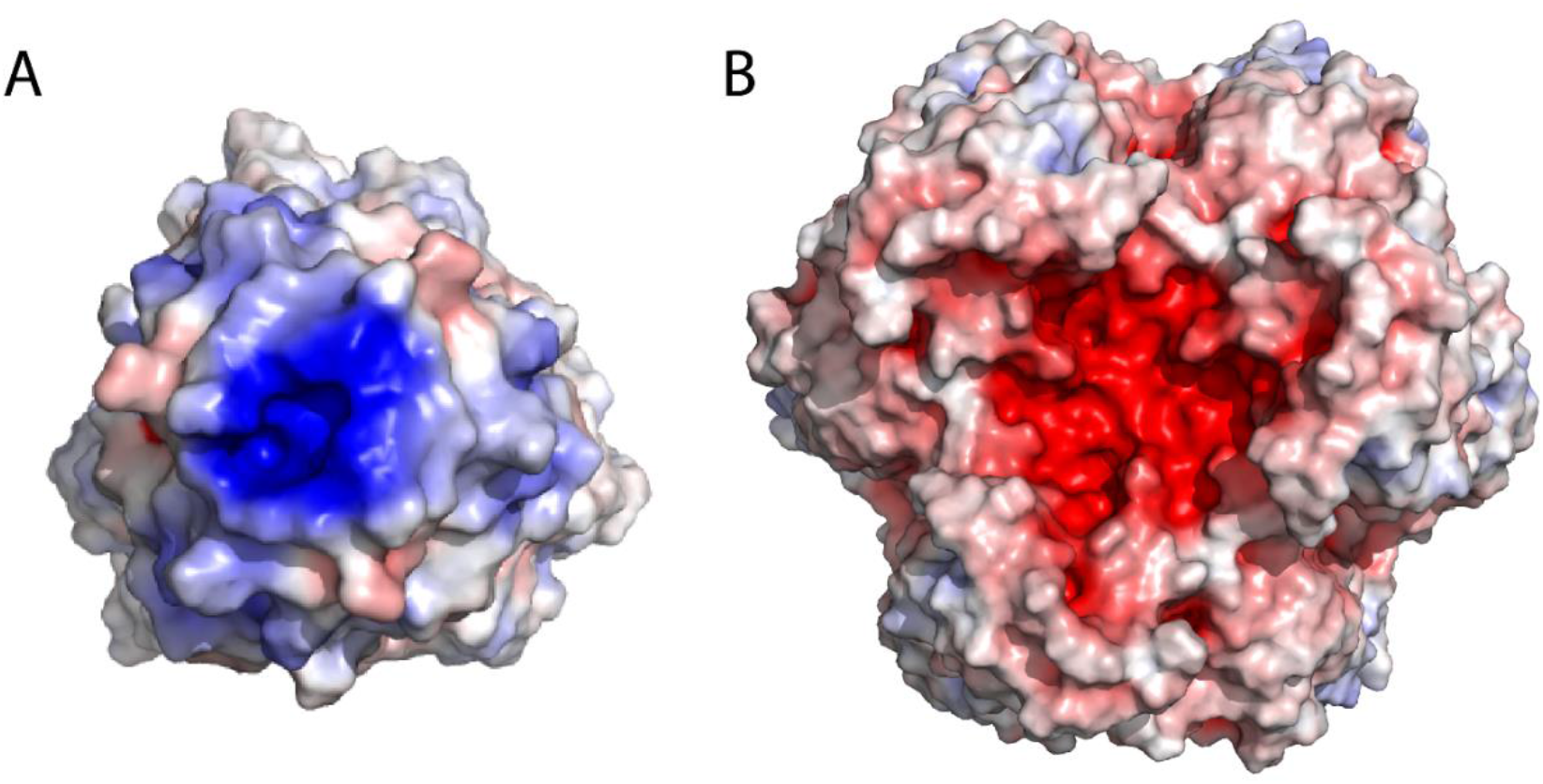
**Surface and charge complementarity** between A] Zav_19 trimer bottom surface and B] AlphaFold (37, 38) Zamilon trimeric MCP (Zav_12) capsomer model, top surface. Electrostatics potential were calculated using APBS (29) and surface colors are fixed at red (– 5 kT/e) to blue (+5 kT/e) and displayed with PyMOL (39).

Interestingly, we found uninterpreted difference density peaks in the vicinity of the β-prism domain, in a pocket at the surface of the prism involving loops from two monomers and lined by polar residues (Thr 22, Asn 23, Asn 31 and Asn 33 – Fig. S1B). This extra density, although not always well defined, is present on all three faces of the prism. It could correspond to a carbohydrate binding site, as described for lectins that also contain a β-prism domain (40–43).

### The Zav_19 fold appears repeated in zamilon

Surprisingly, the Zav_19 β-prism N-terminal domain is conserved in two other proteins of the zamilon virophage, Zav_20 and Zav_1, which show 53% and 51% sequence identity with Zav_19 over 52 amino acids, respectively (Fig. 7A). Zav_20 and Zav_1 are 242 and 309 residues long proteins and, except for the N-terminal domain, share no sequence homology between them or with Zav_19. The central part of the 2 proteins contains repeated G-X-X motifs (14 in Zav_20 and 38 in Zav_1) characteristic of collagen and both proteins are predicted as collagen-like domain containing proteins (Fig. 7A). Interestingly, both C-terminal domains are predicted by AlphaFold (37, 38) to adopt β-sandwich folds (Fig. 7B). Altogether, these results suggest that Zav_20 and Zav_1, as seen for Zav_19, could be trimeric fiber-like proteins. In these proteins, a collagen-like triple-helix structure of different lengths replaces the 3-helix bundle that connects the N-terminal domain to the head domain in Zav_19. We thus hypothesize that Zav_19, Zav_20 and Zav_1 form fibers of different lengths at the surface of the virophage capsid and that the three proteins could be anchored to different MCP capsomers as receptor binding proteins. They could recognize different proteins and/or glycans composing the fibrils of giant viruses as the glycan composition of the different clades can vary significantly (44).

**Figure 7:**
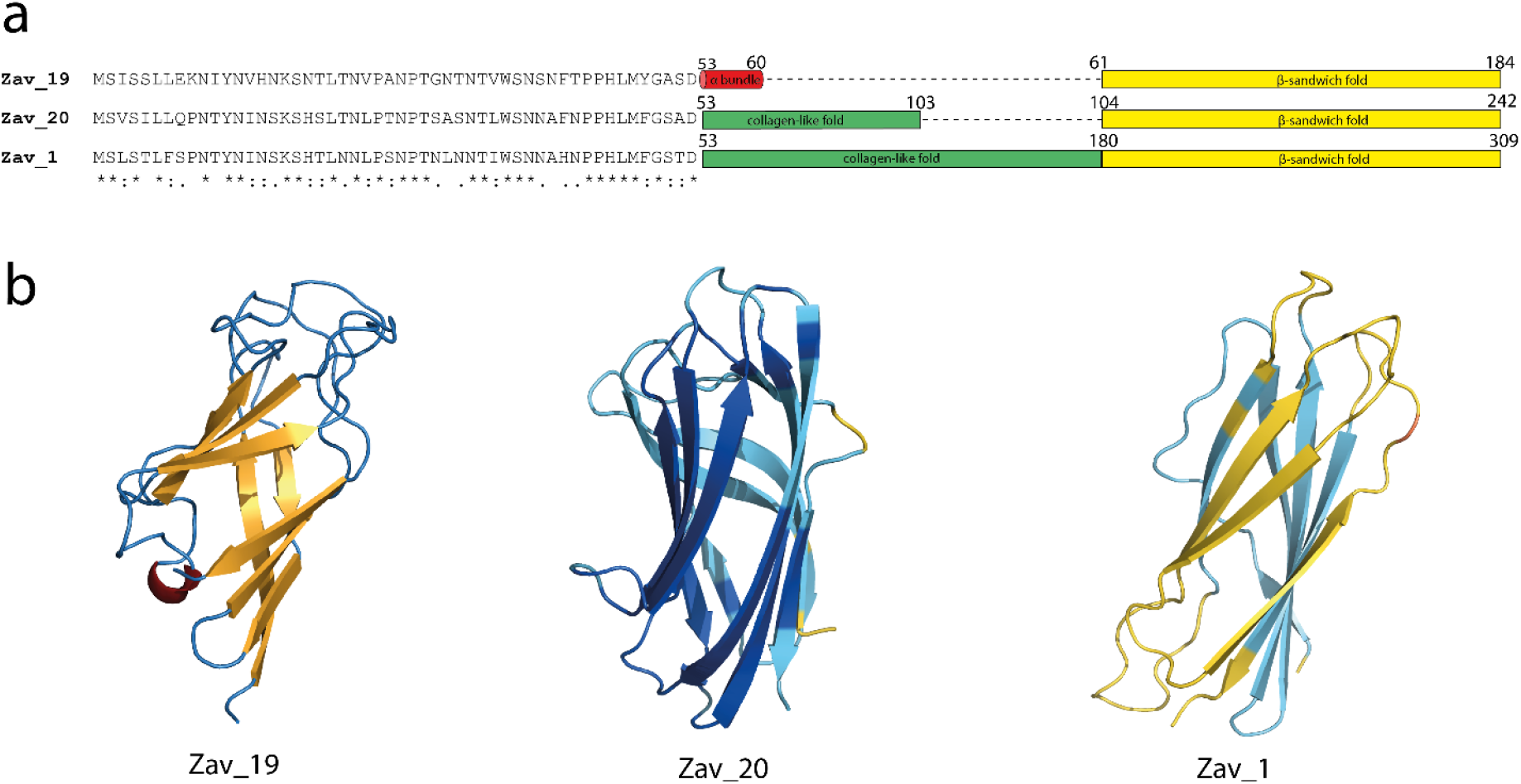
a) Schematic representation of Zav_19, Zav_20 and Zav_1 domains. N-terminal domains were aligned using T-Coffee (45). b) Ribbon representation of the ‘head’ domain of Zav_19 and AlphaFold models of Zav_20 and Zav_1 C-terminal domains. Zav_19 was colored according to secondary structure and Alphafold models were colored by pLDDT score (<50 orange, <70 yellow, <90 light blue, >90 darkblue).

## DISCUSSION

Noumeavirus NMV_189 and zamilon vitis Zav_19 proteins were both predicted as ORFan proteins and are conserved in their own family of viruses, namely *Marseilleviridae* or virophages infecting mimiviruses, respectively. NMV_189 was found to be the most abundant protein in noumeavirus capsids, while Zav_19 ranked 7^th^ in virophage capsids (Table S1). Surprisingly, their 3D-structures revealed a common fold that is also shared with adenoviruses, reoviruses and phages fiber heads of Receptor Binding Proteins (RBP) despite a lack of sequence homology. The noumeavirus protein is, to our knowledge, the first example of an RBP that only contains a head domain, which is supposedly responsible for binding to the host. The surface and charge complementarity between NMV_189 and the noumeavirus MCP model suggested NMV_189 could be nested on top of the capsomers. Their relative abundances in the capsid proteome reinforced this idea, suggesting a one-to-one stoichiometry with the trimeric RBP bound at the center of the trimeric MCP. This was confirmed by recent studies reporting tokyovirus (11) and melbournevirus (31) structures solved at 7.7 and 4.4 Å by cryo-HVEM, where additional density appeared on top of capsomers. The NMV_189 x-ray structure was fitted into the cryo-EM reconstruction, where it perfectly filled the extra density. As a result, NMV_189 increases the capsid diameter by 10 nm, still much lower than the 500 nm required to trigger host phagocytosis. In *Marseilleviridae*, mature neo-synthesized virions exit host cells in giant membrane-limited vesicles, which were proposed to subsequently promote amoeba phagocytosis and trigger the viral replication cycle by an acidification-independent process (46). Virions can also penetrate amoeba cells individually and are found in individual vacuoles, as evidenced by early infection imaging by TEM (12). We thus propose that proteins anchored to MCP capsomers could bind to cell surface receptors and trigger the endocytic process (Fig. S3). The difference map peaks in the binding pocket may correspond to cacodylate molecules in the structure. In a cellular context, since the tokyovirus orthologue of NMV_189 was reported to be glycosylated (11), it could correspond to a glycosylation site involving the virally encoded glycosyltransferases. The glycosylated cap protein NMV_189 could then attach to a glycan receptor of the host cell, triggering phagocytosis and initiation of the infectious cycle. However, since glycosylation of recombinant proteins is not possible in *E. coli*, this hypothesis remains to be tested.

The zamilon vitis fiber head protein Zav_19 is more classical as it is made of two domains, a C-terminal receptor binding domain and an N-terminal β-prism domain, connected by a 3-helix bundle motif. Interestingly, association of a head domain with a β-prism domain was first reported for the gp12 protein of coliphage T4 (47) and for several Lactococcal bacteriophages receptor binding proteins (32, 34, 35, 48). These are also trimeric proteins with three domains, albeit with a different organization: the C-terminal head domain connects to an intermediate β-prism domain, referred to as the neck. The N-terminal domain adopts variable folds in bacteriophages, including a small 3-helix bundle in TP901.1 and Tuc2009 bacteriophages, or a much larger domain that contains α-helices and β-sandwiches, referred to as shoulders in the P2 structure. Virophages that infect *Mimiviridae* are often found associated with giant virus capsids (13, 49, 50), as if they were glued to the heavily glycosylated fibrils surrounding the capsid. It was proposed that this attachment to the fibrils allowed Sputnik and Zamilon to enter host cells with the giant virus (18, 51).

Despite lacking any discernible sequence similarity, lectins can share a β-prism fold and present at least one carbohydrate binding site (40–43). The electron density map of Zav_19 exhibits difference map peaks localized at the periphery of the prism-domain (Fig. S1), which could correspond to carbohydrate binding sites. Intriguingly, the β-prism domain of Zav_19 (residue 1-54) is conserved in 2 other virophage proteins, Zav_20 and Zav_1. As the first structural study of the sputnik virophage capsids structure (36) reported the presence of a mushroom-like fiber with partial occupancy which were not visible in the 3.5 Å resolution map, one possible explanation would be that the capsid of the virophage is covered by the different fiber structures of different lengths that could recognize various ligands, for instance the different sugars composing the fibrils, which are also clade specific (44). The mushroom-like structures could have been the result of averaging in the lower resolution structure and were not resolved, due to their different lengths, in the higher resolution map. These different receptors could be an illustration of the adaptation of virophages to the different glycan compositions of the fibrils of giant viruses they are targeting.

Like the noumeavirus protein, the head domain of Zav_19 is flat due to two long loops involving residue F133 to P144 and K159 to P177 of each monomer and could also bind a receptor on the host cell. In contrast with Giant virus particles, the 250 nm *Marseilleviridae* and 80 nm virophage particles are below the 500 nm threshold needed to trigger Acanthamoeba phagocytosis (1, 52) and could instead use their surface receptor for this purpose. Mavirus virophage infecting the giant virus CroV can penetrate the amoeba host cell in the absence of the giant virus and its genome can be integrated in the cell genome (17). The same could be true for other virophages, even if no integration event in the host cell genome has ever been reported for sputnik or zamilon virophages. It is worth noticing that Zav_1, Zav_19 and Zav_20 are not conserved in mavirus and its host CroV is not covered by a dense layer of fibrils. Thus, the mechanism of entry/interaction could be different in this case.

The receptor binding protein trimer appears as a common fold in the viral world and the conserved head domain can be found associated to various other structural domains. In the case of phages, it was proposed that the RBP head domain could originate from adenoviruses or reoviruses while the other domains (neck and shoulder) could come from gene swapping between viruses (34). As is the case for MCP (53), the similarities between head domains point to a conserved mechanism of action that could be the result of distant evolutionary relationship, either a common origin or lateral gene transfer between different families of viruses. It is probable that other families of icosahedral viruses also use such receptors that have yet to be identified due to their lack of sequence homologies. AlphaFold (37) should help identifying such structures in viral ORFan sequences. It is worth mentioning that African Swine Fever Virus (ASFV) and faustovirus MCPs both exhibit an extra domain forming a crown on top of the capsomers (54, 55). This cap is due to large insertion in the jelly roll domains of the MCP. It is tempting to hypothesize that this supplementary domain could play the role of RBP and be involved in the host recognition. These similarities could also have arisen through convergent evolution as an optimal solution to perform a similar function.

## METHODS

### Genes cloning and protein expression

Genes encoding noumeavirus NMV_189 and zamilon vitis Zav_19 proteins were amplified from noumeavirus and zamilon vitis-containing megavirus vitis genomic DNA, respectively. Genes were inserted by homologous recombination using an In-Fusion HD cloning kit (Clontech) into an in-house modified pETDuet plasmid to yield a cleavable N-terminal His-SUMO fusion protein.

The resulting plasmids were transformed into RIL-LOBSTR strain (56). Cells were grown in 2YT medium containing 100 μg.mL^−1^ ampicillin and 34 μg.mL^−1^ chloramphenicol at 37°C to an A_600_ of 0.7. Temperature was then dropped to 17°C for 20 minutes. Protein expression was induced by adding 0.2 mM of isopropyl β-thiogalactopyranoside, and cells were grown 16– 18 h post induction at 17°C. Bacteria were harvested by centrifugation and resuspended in lysis buffer containing 50 mM Tris-HCl at the appropriate pH (pH 9.0 for NMV_189 and pH 8.0 for Zav_19), 300 mM NaCl, 10 μg.mL^−1^ DNase and EDTA-free protease inhibitor cocktail (Roche). Cells were lysed using sonication and crude lysates were clarified by centrifugation at 13,000 × g for 45 min.

Selenomethionine-substituted proteins were produced using a published protocol to inhibit methionine synthesis in M9 minimal medium in the presence of selenomethionine (57) at 21°C.

### Protein purification

Clarified lysate was applied to a 5 ml HisTrap HP Column (GE Healthcare) equilibrated with buffer A (50 mM Tris-HCl at the appropriate pH, 300 mM NaCl) on an AKTÄ explorer 10S FPLC system (GE Healthcare). The column was washed with 10 column volumes of buffer A, 5 column volumes of buffer A containing 25 mM Imidazole and 5 column volumes of buffer A containing 50 mM Imidazole. Elution was performed with a linear gradient from 50 to 500 m*M* imidazole over 20 column volumes and elution fractions were analyzed by SDS-PAGE.

After cleavage using an in-house His-tagged human rhinovirus 3C protease, a second purification on HisTrap HP was performed to separate the cleaved proteins from the His-Sumo tag and His-tagged protease.

NMV_189 and Zav_19 proteins recovered from the flowthrough were buffer-exchanged using a HiPrep 26/10 column and concentrated to 4.7 mg.mL^−1^ in Tris 10mM pH8.0, NaCl 100mM and 10 mg.mL^−1^ in Tris 10mM pH8.0, respectively. Sample purity was analyzed by SDS-PAGE.

### Nano differential scanning fluorometry

Cacodylate was resuspended in Tris buffer pH 8.0 to a final concentration of 10 mM. Ten μL of pure protein at 10 μM were mixed with 1 μL of cacodylate at 10 mM (ratio 1:100) and placed in nano-DSF grade Prometheus high-sensitivity capillaries and nano-DSF measurements were carried out using a Prometheus NT.48 (Nanotemper technologies GmbH, Munchen, Germany) with a temperature ramp of 1 °C min^−1^, starting at 20 °C and ending at 95 °C with 80% excitation power. Experiments were done in triplicates. Fluorescence data for protein unfolding using fluorescence ratio (F330/F350) was analyzed with the Prometheus software and the melting temperature (Tm) was determined for each sample using the peaks in the first-derivative curve.

### Protein crystallization and structure determination

Recombinant proteins were initially tested at 20°C against 480 different conditions corresponding to commercially available solution sets (SG1, Stura footprint combination HT-96, JCSG+ and wizard screens from Molecular Dimensions) and conditions designed in-house using the *SAmBA* software (58, 59). Screening for crystallization conditions was performed on 3 × 96-well crystallization plates (TPP LabTech) loaded using a *mosquito*^®^ Crystal crystallization robot (TPP Labtech), mixing 0.1 μL protein solution with 0.1 μL reservoir. NMV_189 was crystallized at 20°C by hanging-drop vapor diffusion using 24-well culture plates (Greiner). Each hanging drop was prepared by mixing 0.5 μL NMV_189 (4.7 g.L^−1^) with 0.5 μL of reservoir solution containing 30% PEG 4000 (w/v), 10% MPD (v/v), 0.1 M AmSO4, 0.1 M NaCl and 0.1 M sodium cacodylate pH 6.0 and vapor-equilibrated against 1 mL reservoir solution. Crystals of NMV_189 were flash-frozen in liquid nitrogen. Data collection was carried out at −173°C on the Proxima 1 beamline at the French Synchrotron Soleil (Saclay, France). Diffraction intensities were integrated with XDS (60) and phase determination and structure resolution were performed with the ShelX package (61) implemented into the CCP4 interface (62). Statistics for data collection are listed in Table 1. The 12 possible sites for 12 molecules were found with ShelxD and were used for phasing with ShelxE. The resulting solvent flattened electron density map was used to build the initial SeMet model with Coot (63). Molecular replacement with the SeMet model was used to solve the native structure. The structure was refined against the native data using the Phenix suite (64) at 1.9 Å resolution. The final structure of NMV_189 consists of residues 7 to 153, with 12 molecules per asymmetric unit. Refinement statistics are listed in Table 1. The quality of the model was checked with Molprobity (65). 97.4% of all residues (1713 out 1758) were in the most favored regions of the Ramachandran plot with no outliers.

Zav_19 crystals were obtained by mixing 0.5 μL of reservoir containing 13% to 18% PEG 3350, 0.2 M Ammonium citrate and 0.5 μL of protein concentrated at 10 mg.mL^−1^. Crystals were soaked in 10% glycerol and flash-frozen in liquid nitrogen. Data collection was carried out at −173°C at the Proxima 1 beamline at the French Synchrotron Soleil and the structure was solved using the same strategy as above. Statistics for data collection are listed in Table 1. The 12 possible sites for 6 molecules in the asymmetric unit were found with ShelxD and were used for phasing with ShelxE. The structure was refined at 1.38 Å resolution and consisted of residues 3 to 184. Refinement statistics are listed in Table 1. The quality of the model was checked with Molprobity. 98.0% of all residues (1078 out 1100) were in the most favored regions of the Ramachandran plot and with no outliers.

### Model building and fitting in cryo-electron microscopy maps

The noumeavirus MCP model was built using Phyre2 (66) using the available PBCV-1 MCP VP54 structure (28) as template (PDB: 5TIP). Fitting of the NMV_189 structure in the melbournevirus cryo-EM map was performed with ChimeraX (67).

## ACKNOWLEDGMENTS

This work was partially supported by the French National Research Agency ANR-20-CE11-0001, Institut Français de Bioinformatique (ANR–11–INSB–0013), the Fondation Bettencourt-Schueller (OTP51251), the Provence-Alpes-Côte-d’Azur région (2010 12125), and the Excellence Initiative of Aix-Marseille Université – A*MIDEX, a french “Investissements d’Avenir” program AMX-18-ACE-004. Proteomic experiments were partly supported by the ProFi grant (ANR-10-INBS-08-01). We also acknowledge the support of the bottom-up platform and informatics group of EDyP, Pierre Legrand for his help solving structures on Proxima 1 beamline at SOLEIL, Kazuyoshi Murata and Kenta Okamoto for providing Cryo-EM map prior public release and Carla Rossi for technical help.

## AUTHOR CONTRIBUTIONS

S.J. and C.A. conceptualization, methodology, validation, formal analysis, investigation, visualization, writing – original draft; E.G. methodology, investigation, writing – review & visualization; A.S. methodology, software, investigation

## DATA AVAILABILITY

NMV_189 and Zav_19 coordinates and structure factors have been deposited in the Protein Data Bank with accession number 7QRR and 7QRJ, respectively.

## SUPPLEMENTARY INFORMATION

**Figure S1:**
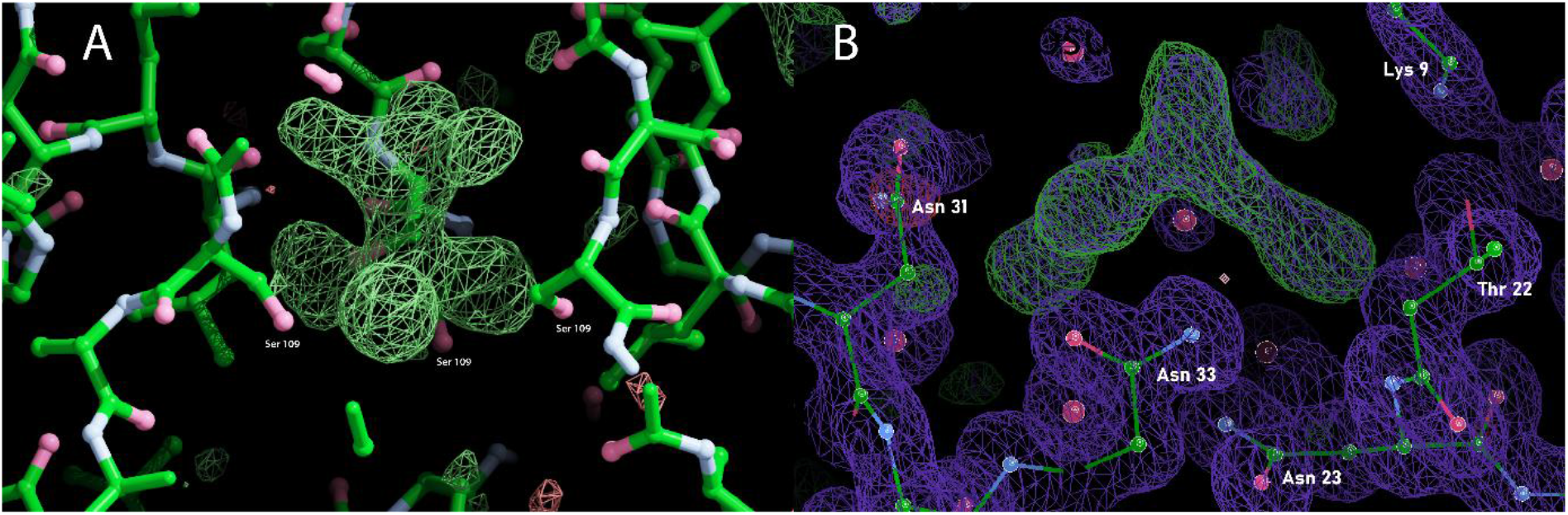
**Difference map** showing an unmodelled blob of density A) in the channel of the NMV_189 trimeric structure B) at the periphery of the β-prism domain of Zav_19.

**Figure S2:**
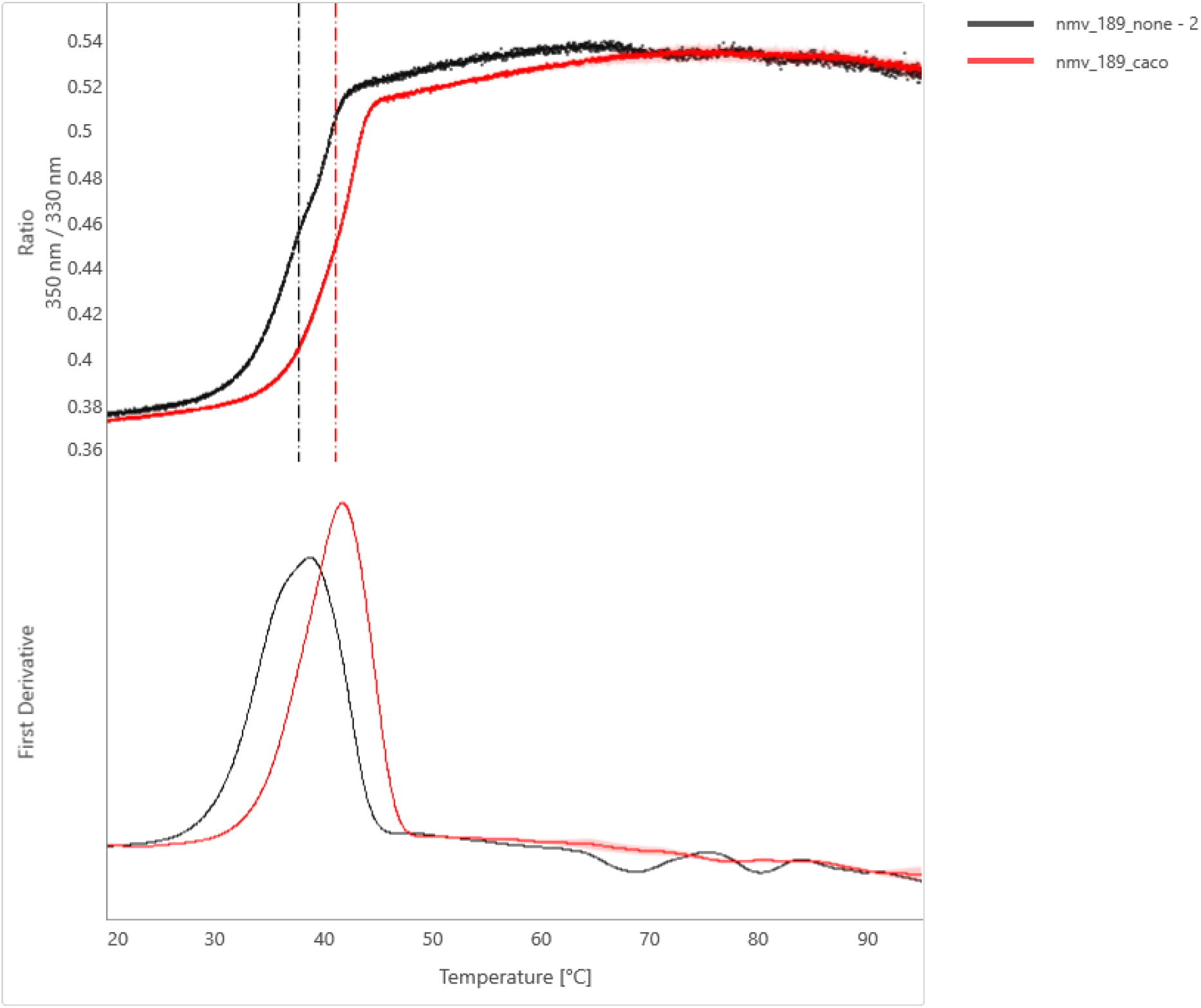
NanoDSF curves (top) and their first derivative (bottom) of NMV_189 alone (black line) or in presence of cacodylate (red line).

**Figure S3:**
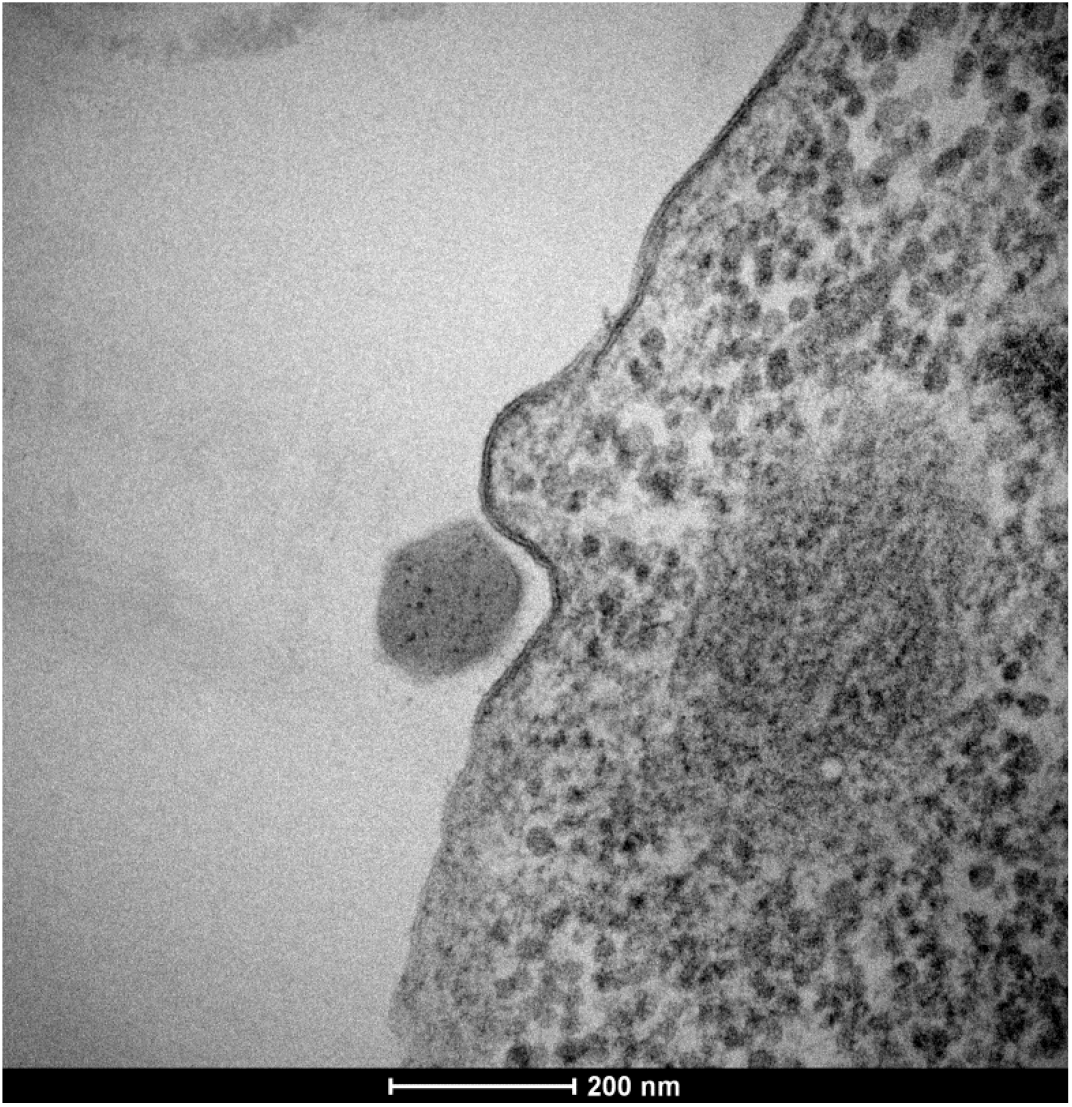
**Transmission Electron micrograph** of a single particle of Noumeavirus triggering Acanthamoeba cell endocytosis.

**Table S1:** **Ranking of the most abundant proteins** identified in Noumeavirus and Zamilon vitis purified particles by mass spectroscopy

| Noumeavirus proteins |  |  | Zamilon vitis proteins |  |  |
| --- | --- | --- | --- | --- | --- |
| Protein Ids | Annotation | Rank | Protein Ids | Annotation | Rank |
| NMV_189 | hypothetical protein | 1 | Zav_12 | Major Capsid protein | 1 |
| NMV_078 | histone H3-like protein | 2 | Zav_13 | Endonuclease DNA binding | 2 |
| NMV_116 | major capsid protein | 3 | Zav_11 | Minor Capsid protein | 3 |
| NMV_149 | hypothetical protein | 4 | Zav_4 | DNA packaging ATPase | 4 |
| NMV_171 | papain-like cysteine protease | 5 | Zav_6 | Putative DNA binding | 5 |
| NMV_079 | histone H2B/H2A fusion protein | 6 | Zav_18 | Cysteine protease | 6 |
| NMV_185 | hypothetical protein | 7 | Zav_19 | Hypothetical | 7 |
| NMV_333 | hypothetical protein | 8 | Mvi_320 | Putative DNA binding | 8 |
| NMV_141 | hypothetical protein | 9 | Zav_20 | Putative collagen domain | 9 |
| NMV_192 | hypothetical protein | 10 | Mvtv_7 | Zinc Finger domain | 10 |
| NMV_225 | thioredoxin | 11 | Zav_1 | Collagen-like protein | 11 |
| NMV_230 | hypothetical protein | 12 | Mvi_621 | Putative DNA binding | 12 |
| NMV_045 | hypothetical protein | 13 | g557-519644-520579 | Vtype ATPase, C subunit; membrane protein | 13 |
| NMV_238 | hypothetical protein | 14 | Mvi_114 | Hypothetical | 14 |
| NMV_066 | hypothetical protein | 15 | Zav_5 | Putative DNA binding | 15 |
| NMV_349 | hypothetical protein | 16 | Zav_17 | Putative integrase | 16 |
| NMV_163 | hypothetical protein | 17 | g641-1372693-1373537 | Putative signal peptide | 17 |
| NMV_198 | membrane protein | 18 | Mvi_472 | Ubiquitin-like protein | 18 |
| NMV_068 | AAA-family ATPase | 19 | Zav_10 | Putative transposase | 19 |
| NMV_296 | hypothetical protein | 20 | Zav_14 | Putative Zing Finger C2H2; DNA binding | 20 |
| NMV_058 | galactose binding lectin domain containing protein | 21 | Zav_3 | Putative Zing Finger C2H2; DNA binding | 21 |
| NMV_060 | transmembrane domain containing protein | 22 | g254-27747-28588 | Putative signal peptide | 22 |
| NMV_183 | membrane protein | 23 | Mvi_125 | Putative DNA binding | 23 |
| NMV_350 | hypothetical protein | 24 | Mvi_142 | Putative signal peptide | 24 |
| NMV_355 | transmembrane domain containing protein | 25 | Mvi_825 | Putative GMC-type oxidoreductase; membrane | 25 |

## REFERENCES

1. Boyer M, Yutin N, Pagnier I, Barrassi L, Fournous G, Espinosa L, Robert C, Azza S, Sun S, Rossmann MG, Suzan-Monti M, La Scola B, Koonin EV, Raoult D. 2009. Giant Marseillevirus highlights the role of amoebae as a melting pot in emergence of chimeric microorganisms. Proc Natl Acad Sci U S A 106:21848–21853.

2. Blanca L, Christo-Foroux E, Rigou S, Legendre M. 2020. Comparative Analysis of the Circular and Highly Asymmetrical Marseilleviridae Genomes. Viruses 12:1270.

3. Thomas V, Bertelli C, Collyn F, Casson N, Telenti A, Goesmann A, Croxatto A, Greub G. 2011. Lausannevirus, a giant amoebal virus encoding histone doublets. Environ Microbiol 13:1454–1466.

4. Aherfi S, Boughalmi M, Pagnier I, Fournous G, La Scola B, Raoult D, Colson P. 2014. Complete genome sequence of Tunisvirus, a new member of the proposed family Marseilleviridae. Arch Virol 159:2349–2358.

5. Dornas FP, Assis FL, Aherfi S, Arantes T, Abrahão JS, Colson P, La Scola B. 2016. A Brazilian Marseillevirus Is the Founding Member of a Lineage in Family Marseilleviridae. Viruses 8:76.

6. Dos Santos RN, Campos FS, Medeiros de Albuquerque NR, Finoketti F, Côrrea RA, Cano-Ortiz L, Assis FL, Arantes TS, Roehe PM, Franco AC. 2016. A new marseillevirus isolated in Southern Brazil from Limnoperna fortunei. Sci Rep 6:35237.

7. Doutre G, Philippe N, Abergel C, Claverie J-M. 2014. Genome Analysis of the First Marseilleviridae Representative from Australia Indicates that Most of Its Genes Contribute to Virus Fitness. J Virol 88:14340–14349.

8. Doutre G, Arfib B, Rochette P, Claverie J-M, Bonin P, Abergel C. 2015. Complete Genome Sequence of a New Member of the Marseilleviridae Recovered from the Brackish Submarine Spring in the Cassis Port-Miou Calanque, France. Genome Announc 3.

9. Aherfi S, Pagnier I, Fournous G, Raoult D, La Scola B, Colson P. 2013. Complete genome sequence of Cannes 8 virus, a new member of the proposed family “Marseilleviridae.” Virus Genes 47:550–555.

10. Okamoto K, Miyazaki N, Reddy HKN, Hantke MF, Maia FRNC, Larsson DSD, Abergel C, Claverie J-M, Hajdu J, Murata K, Svenda M. 2018. Cryo-EM structure of a Marseilleviridae virus particle reveals a large internal microassembly. Virology 516:239–245.

11. Chihara A, Burton-Smith RN, Kajimura N, Mitsuoka K, Okamoto K, Song C, Murata K. 2021. A novel capsid protein network allows the characteristic inner membrane structure of Marseilleviridae giant viruses. bioRxiv 2021.02.03.428533.

12. Fabre E, Jeudy S, Santini S, Legendre M, Trauchessec M, Couté Y, Claverie J-M, Abergel C. 2017. Noumeavirus replication relies on a transient remote control of the host nucleus. Nat Commun 8.

13. La Scola, B. et al. The virophage as a unique parasite of the giant mimivirus. Nature 455, 100–104 (2008).

14. Mougari S, Bekliz M, Abrahao J, Di Pinto F, Levasseur A, La Scola B. 2019. Guarani Virophage, a New Sputnik-Like Isolate From a Brazilian Lake. Front Microbiol 10:1003.

15. Fischer MG, Allen MJ, Wilson WH, Suttle CA. 2010. Giant virus with a remarkable complement of genes infects marine zooplankton. Proc Natl Acad Sci U S A 107:19508–19513.

16. Fischer MG, Suttle CA. 2011. A Virophage at the Origin of Large DNA Transposons. Science 332:231–234.

17. Fischer MG, Hackl T. 2016. Host genome integration and giant virus-induced reactivation of the virophage mavirus. Nature 540:288–291.

18. Gaia M, Benamar S, Boughalmi M, Pagnier I, Croce O, Colson P, Raoult D, La Scola B. 2014. Zamilon, a Novel Virophage with Mimiviridae Host Specificity. PLoS ONE 9.

19. Jeudy S, Bertaux L, Alempic J-M, Lartigue A, Legendre M, Belmudes L, Santini S, Philippe N, Beucher L, Biondi EG, Juul S, Turner DJ, Couté Y, Claverie J-M, Abergel C. 2019. Exploration of the propagation of transpovirons within Mimiviridae reveals a unique example of commensalism in the viral world. ISME J 1–13.

20. Desnues C, La Scola B, Yutin N, Fournous G, Robert C, Azza S, Jardot P, Monteil S, Campocasso A, Koonin EV, Raoult D. 2012. Provirophages and transpovirons as the diverse mobilome of giant viruses. Proc Natl Acad Sci U S A 109:18078–18083.

21. Zhang X, Sun S, Xiang Y, Wong J, Klose T, Raoult D, Rossmann MG. 2012. Structure of Sputnik, a virophage, at 3.5-Å resolution. Proc Natl Acad Sci U S A 109:18431–18436.

22. NCBI Resource Coordinators. 2016. Database resources of the National Center for Biotechnology Information. Nucleic Acids Res 44:D7–D19.

23. Holm L, Laakso LM. 2016. Dali server update. Nucleic Acids Res 44:W351–355.

24. Gibrat JF, Madej T, Bryant SH. 1996. Surprising similarities in structure comparison. Curr Opin Struct Biol 6:377–385.

25. Guardado-Calvo P, Llamas-Saiz AL, Fox GC, Langlois P, van Raaij MJ. 2007. Structure of the C-terminal head domain of the fowl adenovirus type 1 long fiber. J Gen Virol 88:2407–2416.

26. El Bakkouri M, Seiradake E, Cusack S, Ruigrok RWH, Schoehn G. 2008. Structure of the C-terminal head domain of the fowl adenovirus type 1 short fibre. Virology 378:169–176.

27. Broeker NK, Roske Y, Valleriani A, Stephan MS, Andres D, Koetz J, Heinemann U, Barbirz S. 2019. Time-resolved DNA release from an O-antigen-specific Salmonella bacteriophage with a contractile tail. J Biol Chem 294:11751–11761.

28. De Castro C, Klose T, Speciale I, Lanzetta R, Molinaro A, Van Etten JL, Rossmann MG. 2018. Structure of the chlorovirus PBCV-1 major capsid glycoprotein determined by combining crystallographic and carbohydrate molecular modeling approaches. Proc Natl Acad Sci U S A 115:E44–E52.

29. Baker NA, Sept D, Joseph S, Holst MJ, McCammon JA. 2001. Electrostatics of nanosystems: Application to microtubules and the ribosome. Proc Natl Acad Sci U S A 98:10037–10041.

30. Pettersen EF, Goddard TD, Huang CC, Couch GS, Greenblatt DM, Meng EC, Ferrin TE. 2004. UCSF Chimera-A visualization system for exploratory research and analysis. J Comput Chem 25:1605–1612.

31. Burton-Smith RN, Reddy HKN, Svenda M, Abergel C, Okamoto K, Murata K. 2021. The 4.4 Å structure of the giant Melbournevirus virion belonging to the Marseilleviridae family. bioRxiv https://doi.org/10.1101/2021.07.14.452405.

32. Bebeacua C, Bron P, Lai L, Vegge CS, Brøndsted L, Spinelli S, Campanacci V, Veesler D, Heel M van, Cambillau C. 2010. Structure and Molecular Assignment of Lactococcal Phage TP901-1 Baseplate *. J Biol Chem 285:39079–39086.

33. Spinelli S, Campanacci V, Blangy S, Moineau S, Tegoni M, Cambillau C. 2006. Modular Structure of the Receptor Binding Proteins of Lactococcus lactis Phages: THE RBP STRUCTURE OF THE TEMPERATE PHAGE TP901-1 *. J Biol Chem 281:14256–14262.

34. Spinelli S, Desmyter A, Verrips CT, de Haard HJW, Moineau S, Cambillau C. 2006. Lactococcal bacteriophage p2 receptor-binding protein structure suggests a common ancestor gene with bacterial and mammalian viruses. 1. Nat Struct Mol Biol 13:85–89.

35. Legrand P, Collins B, Blangy S, Murphy J, Spinelli S, Gutierrez C, Richet N, Kellenberger C, Desmyter A, Mahony J, van Sinderen D, Cambillau C. 2016. The Atomic Structure of the Phage Tuc2009 Baseplate Tripod Suggests that Host Recognition Involves Two Different Carbohydrate Binding Modules. mBio 7:e01781–15.

36. Sun S, La Scola B, Bowman VD, Ryan CM, Whitelegge JP, Raoult D, Rossmann MG. 2010. Structural Studies of the Sputnik Virophage. J Virol 84:894–897.

37. Jumper J, Evans R, Pritzel A, Green T, Figurnov M, Ronneberger O, Tunyasuvunakool K, Bates R, Žídek A, Potapenko A, Bridgland A, Meyer C, Kohl SAA, Ballard AJ, Cowie A, Romera-Paredes B, Nikolov S, Jain R, Adler J, Back T, Petersen S, Reiman D, Clancy E, Zielinski M, Steinegger M, Pacholska M, Berghammer T, Bodenstein S, Silver D, Vinyals O, Senior AW, Kavukcuoglu K, Kohli P, Hassabis D. 2021. Highly accurate protein structure prediction with AlphaFold. Nature 596:583–589.

38. Varadi M, Anyango S, Deshpande M, Nair S, Natassia C, Yordanova G, Yuan D, Stroe O, Wood G, Laydon A, Žídek A, Green T, Tunyasuvunakool K, Petersen S, Jumper J, Clancy E, Green R, Vora A, Lutfi M, Figurnov M, Cowie A, Hobbs N, Kohli P, Kleywegt G, Birney E, Hassabis D, Velankar S. 2022. AlphaFold Protein Structure Database: massively expanding the structural coverage of protein-sequence space with high-accuracy models. Nucleic Acids Res 50:D439–D444.

39. Schrödinger L,, DeLano W. 2020. PyMOL. http://www.pymol.org/pymol.

40. Sharma A, Chandran D, Singh DD, Vijayan M. 2007. Multiplicity of carbohydrate-binding sites in beta-prism fold lectins: occurrence and possible evolutionary implications. J Biosci 32:1089–1110.

41. Cabanettes A, Perkams L, Spies C, Unverzagt C, Varrot A. 2018. Recognition of Complex Core-Fucosylated N-Glycans by a Mini Lectin. Angew Chem Int Ed Engl 57:10178–10181.

42. Yamasaki K, Kubota T, Yamasaki T, Nagashima I, Shimizu H, Terada R-I, Nishigami H, Kang J, Tateno M, Tateno H. 2019. Structural basis for specific recognition of core fucosylation in N-glycans by Pholiota squarrosa lectin (PhoSL). Glycobiology 29:576–587.

43. Yamasaki K, Yamasaki T, Tateno H. 2018. The trimeric solution structure and fucose-binding mechanism of the core fucosylation-specific lectin PhoSL. 1. Sci Rep 8:7740.

44. Notaro A, Poirot O, Garcin ED, Nin S, Molinaro A, Tonetti M, De Castro C, Abergel C. 2022. Giant viruses of the Megavirinae subfamily possess biosynthetic pathways to produce rare bacterial-like sugars in a clade-specific manner. microLife 3:uqac002.

45. Notredame C, Higgins DG, Heringa J. 2000. T-coffee: a novel method for fast and accurate multiple sequence alignment11Edited by J. Thornton. J Mol Biol 302:205–217.

46. Arantes TS, Rodrigues RAL, Dos Santos Silva LK, Oliveira GP, de Souza HL, Khalil JYB, de Oliveira DB, Torres AA, da Silva LL, Colson P, Kroon EG, da Fonseca FG, Bonjardim CA, La Scola B, Abrahão JS. 2016. The Large Marseillevirus Explores Different Entry Pathways by Forming Giant Infectious Vesicles. J Virol 90:5246–5255.

47. van Raaij MJ, Schoehn G, Burda MR, Miller S. 2001. Crystal structure of a heat and protease-stable part of the bacteriophage T4 short tail fibre. J Mol Biol 314:1137–1146.

48. Veesler D, Spinelli S, Mahony J, Lichière J, Blangy S, Bricogne G, Legrand P, Ortiz-Lombardia M, Campanacci V, Sinderen D van, Cambillau C. 2012. Structure of the phage TP901-1 1.8 MDa baseplate suggests an alternative host adhesion mechanism. Proc Natl Acad Sci 109:8954–8958.

49. Desnues C, Raoult D. 2010. Inside the Lifestyle of the Virophage. Intervirology 53:293–303.

50. Duponchel S, Fischer MG. 2019. Viva lavidaviruses! Five features of virophages that parasitize giant DNA viruses. PLoS Pathog 15.

51. Taylor BP, Cortez MH, Weitz JS. 2014. The virus of my virus is my friend: Ecological effects of virophage with alternative modes of coinfection. J Theor Biol 354:124–136.

52. Korn ED, Weisman RA. 1967. Phagocytosis of latex beads by acanthamoeba. J Cell Biol 34:219–227.

53. Benson SD, Bamford JK, Bamford DH, Burnett RM. 1999. Viral evolution revealed by bacteriophage PRD1 and human adenovirus coat protein structures. Cell 98:825–833.

54. Klose T, Reteno DG, Benamar S, Hollerbach A, Colson P, La Scola B, Rossmann MG. 2016. Structure of faustovirus, a large dsDNA virus. Proc Natl Acad Sci U S A 113:6206–6211.

55. Andrés G, Charro D, Matamoros T, Dillard RS, Abrescia NGA. 2020. The cryo-EM structure of African swine fever virus unravels a unique architecture comprising two icosahedral protein capsids and two lipoprotein membranes. J Biol Chem 295:1–12.

56. Andersen KR, Leksa NC, Schwartz TU. 2013. Optimized E. coli expression strain LOBSTR eliminates common contaminants from His-tag purification. Proteins 81:1857–1861.

57. Doublié S. 1997. [29] Preparation of selenomethionyl proteins for phase determination, p. 523–530. *In* Methods in Enzymology. Academic Press.

58. Abergel C, Moulard M, Moreau H, Loret E, Cambillau C, Fontecilla-Camps JC. 1991. Systematic use of the incomplete factorial approach in the design of protein crystallization experiments. J Biol Chem 266:20131–20138.

59. Audic S, Lopez F, Claverie JM, Poirot O, Abergel C. 1997. SAmBA: an interactive software for optimizing the design of biological macromolecules crystallization experiments. Proteins 29:252–257.

60. Kabsch W. 2010. XDS. Acta Crystallogr D Biol Crystallogr 66:125–132.

61. Sheldrick GM. 2008. A short history of SHELX. 1. Acta Crystallogr A 64:112–122.

62. Winn MD, Ballard CC, Cowtan KD, Dodson EJ, Emsley P, Evans PR, Keegan RM, Krissinel EB, Leslie AGW, McCoy A, McNicholas SJ, Murshudov GN, Pannu NS, Potterton EA, Powell HR, Read RJ, Vagin A, Wilson KS. 2011. Overview of the CCP4 suite and current developments. 4. Acta Crystallogr D Biol Crystallogr 67:235–242.

63. Emsley P, Lohkamp B, Scott WG, Cowtan K. 2010. Features and development of *Coot*. Acta Crystallogr D Biol Crystallogr 66:486–501.

64. Liebschner D, Afonine PV, Baker ML, Bunkóczi G, Chen VB, Croll TI, Hintze B, Hung L-W, Jain S, McCoy AJ, Moriarty NW, Oeffner RD, Poon BK, Prisant MG, Read RJ, Richardson JS, Richardson DC, Sammito MD, Sobolev OV, Stockwell DH, Terwilliger TC, Urzhumtsev AG, Videau LL, Williams CJ, Adams PD. 2019. Macromolecular structure determination using X-rays, neutrons and electrons: recent developments in Phenix. 10. Acta Crystallogr Sect Struct Biol 75:861–877.

65. Williams CJ, Headd JJ, Moriarty NW, Prisant MG, Videau LL, Deis LN, Verma V, Keedy DA, Hintze BJ, Chen VB, Jain S, Lewis SM, Arendall WB, Snoeyink J, Adams PD, Lovell SC, Richardson JS, Richardson DC. 2017. MolProbity: More and better reference data for improved all-atom structure validation. Protein Sci 27:293–315.

66. Kelley LA, Mezulis S, Yates CM, Wass MN, Sternberg MJE. 2015. The Phyre2 web portal for protein modeling, prediction and analysis. Nat Protoc 10:845–858.

67. Pettersen EF, Goddard TD, Huang CC, Meng EC, Couch GS, Croll TI, Morris JH, Ferrin TE. 2021. UCSF ChimeraX: Structure visualization for researchers, educators, and developers. Protein Sci Publ Protein Soc 30:70–82.

## References

1. Fabre E, Jeudy S, Santini S, Legendre M, Trauchessec M, Couté Y, Claverie J-M, Abergel C. 2017. Noumeavirus replication relies on a transient remote control of the host nucleus. Nat Commun 8:15087.

2. Jeudy S, Bertaux L, Alempic J-M, Lartigue A, Legendre M, Belmudes L, Santini S, Philippe N, Beucher L, Biondi EG, Juul S, Turner DJ, Couté Y, Claverie J-M, Abergel C. 2019. Exploration of the propagation of transpovirons within Mimiviridae reveals a unique example of commensalism in the viral world. ISME J 1–13.

